# MotifPeeker: Benchmarking epigenomic profiling methods using motifs

**DOI:** 10.1101/2025.03.31.645756

**Authors:** Hiranyamaya Dash, Thomas Roberts, Maria Weinert, Nathan Skene

## Abstract

MotifPeeker benchmarks epigenomic profiling methods targeting transcription factors (TFs) where no “gold standard” reference exists, using motif enrichment as a key metric. With minimal input, users can analyse their data in a single function and receive an intuitive HTML report.

**Availability and Implementation:** MotifPeeker is available on Bioconductor (≥ v3.21) at https://bioconductor.org/packages/MotifPeeker. The complete source code is available on GitHub at https://github.com/neurogenomics/MotifPeeker, with full documentation provided at https://neurogenomics.github.io/MotifPeeker. Additionally, the MotifPeeker Docker image is hosted on GitHub at https://github.com/neurogenomics/MotifPeeker/pkgs/container/motifpeeker.

## Introduction

Understanding the cell type-specific epigenome has become vital for understanding the role of genetic variants linked to common diseases because the majority of disease-associated SNPs appear to affect non-coding regulatory regions such as promoters and enhancers (Finucane et al., 2015). The majority of the epigenetic data collected to date has been from easily accessible bulk samples (e.g. whole blood) or cell lines, however, because the epigenome varies greatly between cell types, and it is these cell-type specific features which are most relevant to disease, there has been a strong push in recent years to develop new methods better suited for profiling single cell types from heterogenous tissues. The classical technique used in the field has been Chromatin immunoprecipitation followed by sequencing (ChIP-Seq): it is now a well-established technique for performing genome-wide mapping of regulatory protein-DNA interactions (Dasgupta and Chellappan, 2007; Gade and Kalvakolanu, 2012), however, it requires large numbers of cells and generates a high background signal. Recent techniques, such as Cleavage under targets and tagmentation (CUT&Tag) (Kaya-Okur et al., 2019) and Targeted insertion of promoters sequencing (TIP-Seq) (Bartlett et al., 2021), utilise Tn5 transposase activity to perform more precise, targeted cuts, improving the sensitivity and resolution of the experiments, making it possible to profile proteins in as few as a single cell (Bartosovic et al., 2021).

However, the validation of recently developed epigenomic profiling methods against established techniques such as ChIP-Seq remains under-explored, particularly regarding their data generation fidelity and the robustness of their analytical frameworks. A typical scenario to consider is when a new method, owing to its greater sensitivity, reports many more candidate sites than ChIP Seq. This prompts the question of whether these extra peaks are genuine transcription factor binding sites or artefacts of increased detection. While most studies, including the original papers that introduced these assays, have benchmarked their results against traditional ChIP-Seq, little systematic work has been done to validate peaks unique to the newer methods (Abbasova et al., 2025).

In addition to the variations in output between different epigenomic profiling methods, significant differences can also arise from the same method, depending on the specific experimental protocol and data processing strategy used. Tools such as ChIPComp (Chen et al., 2015) and DiffBind (Stark and Brown, 2017) allow users to perform differential binding analysis of ChIP-seq data, identifying genomic regions with significant differences in protein–DNA binding between conditions. These methods are valuable within a single assay type but offer little for cross-method bench-marking, as they rely on comparing read counts at shared peaks, an approach that presumes similar signal and noise properties across datasets, conditions met within ChIP-Seq but not when contrasting different assays. EpiCompare (Choi et al., 2023) begins to address systematic comparisons across different profiling methods by stringing together a range of downstream analyses that were previously spread across various software, yet no ideal strategy has been established for such comparisons. Moreover, EpiCompare requires a dataset to be defined as a “gold standard”: with the implicit assumption that an ENCODE ChIP-Seq dataset would be used for this purpose. There is, however, no intrinsic reason to believe that ChIP-Seq is a more faithful representation of epigenetic profiles than that which can be obtained with newer methods.

To mitigate the need of a gold standard dataset, we propose instead to use motif enrichment as the criterion for assessing and comparing epigenomic data. DNA motifs are short, recurring sequences believed to serve a biological purpose. They frequently mark the binding sites for proteins such as transcription factors (TFs) that regulate gene expression (D’haeseleer, 2006). Previous studies have established the presence of highly enriched motifs in most ChIP-Seq data, revealing both new motifs and validating known ones (Kheradpour and Kellis, 2014; Wang et al., 2012). Regions of high signals within epigenomic data (peaks) can be considered more legitimate if they contain a higher proportion of motifs relevant to the target protein. Our R package, MotifPeeker, generates an intuitive and descriptive report file comparing user-provided peak files using our proposed approach. It does not require the reference dataset to be inherently better; it simply aids in evaluating the relative performances of experiments. The ability to run the entire analysis with a single function call from the package makes our approach more accessible and time-saving for users with limited computational experience. While existing tools such as MEME-ChIP (Ma et al., 2014) perform motif discovery and enrichment analysis within individual datasets, MotifPeeker extends this by providing a comparative framework, enabling evaluation across multiple experiments.

## Implementation

MotifPeeker was implemented using R programming language version 4.4.1 (R Core Team, 2024). The code was written in compliance with the Bioconductor coding and documentation standards (Gentleman et al., 2004). We used rworkflows as our package’s continuous integration (CI) setup solution to automatically deploy the documentation website and run all unit tests in the cloud to maintain the integrity of the package functions (Schilder et al., 2024). The entire analysis can be executed with a single function, *MotifPeeker*, which shares its name with the package. The principal input required for a MotifPeeker run is the peak files, with support currently for files generated by MACS and SEACR (Meers et al., 2019; Zhang et al., 2008). The input files are expected to correspond to experiments targeting the same protein and using the same genome build. MotifPeeker is therefore designed for datasets profiling the same transcription factor across methods. It is not suitable for broader assays such as ATAC-Seq, which do not target a single factor. The function parses the parameters, passes them into an RMarkdown file, and calls helper functions to render the final HTML report, as illustrated in Figure 1.

**Figure 1.**
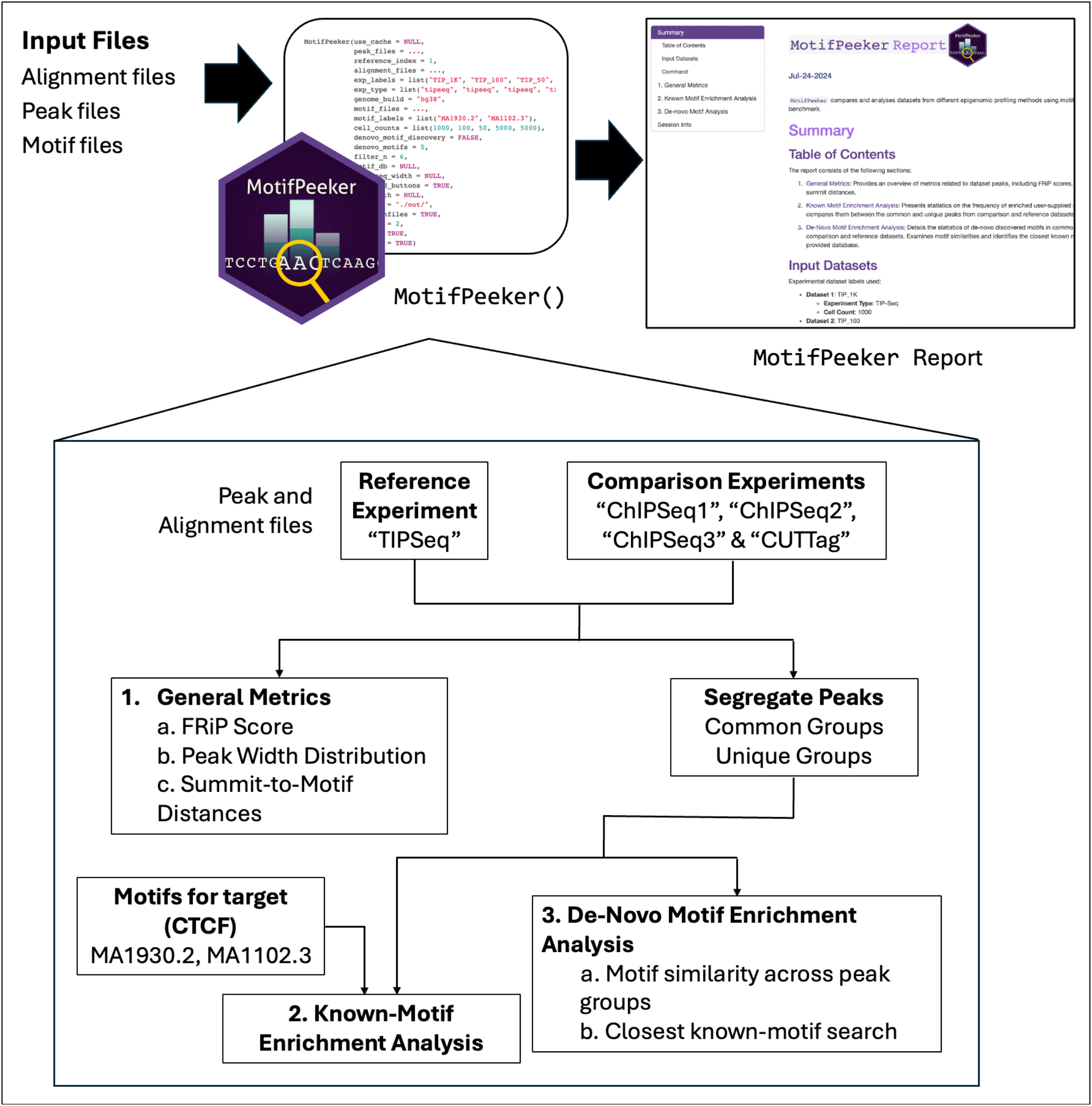
Workflow for the MotifPeeker package. Users only need to call the MotifPeeker function with all the input files supplied. Various parameters, such as the reference dataset and number of motifs to discover, can be adjusted for optional customizability. Once the run is complete, an HTML report is opened, which guides the viewer through all the analysis plots. The three sections of the reports are numbered, with their outputs described in detail in Figure 2.

The analysis is divided into three sections: (1) General Statistics, which provide an overview of metrics related to dataset peaks; (2) Known Motif Enrichment Analysis, which presents motif enrichment counts for the user-supplied motifs; and (3) Discovered Motif Analysis, which examines the similarities of motifs discovered in the comparisons and identifies their closest known matches in the JASPAR database (Rauluseviciute et al., 2024). For sections 2 and 3, a pairwise comparison is conducted between each additional peak file and the first. In each pairwise comparison, the peaks from both experiments are divided into four groups based on their overlap, using the *findOverlaps* function from the GenomicRanges package (Lawrence et al., 2013). This segregation of peaks into common and unique groups allows assessment of whether either group captures a higher number of motifs, indicated by relatively higher enrichment in the unique peak group.

MotifPeeker uses the MEME Suite—an established toolkit for motif analysis since 1994—through memes, its R interface (Bailey et al., 2015; Nystrom and McKay, 2021). This suite includes several tools for motif analysis, including AME (Analysis of Motif Enrichment), FIMO (Find Individual Motif Occurrences), STREME (Simple, Thorough, Rapid, Enriched Motif Elicitation) and TOMTOM. AME is designed to detect the enrichment of known, user-provided motifs in a set of sequences. FIMO finds all the true occurrences of user-provided motifs in the input sequences by comparing the likelihood of each possible occurrence of the motifs in the input to the background set of sequences (Grant et al., 2011). For motif discovery, STREME generates suffix trees to count all the k-mer possibilities in the input and control sequences, followed by checking for relative enrichment of each k-mer compared to the background sequences (Bailey, 2021). Lastly, TOMTOM is used to find the closest known motif to the input sequences in a supplied database of known motifs, such as JASPAR (Gupta et al., 2007). Computationally intensive tools, particularly STREME, are multithreaded using BiocParallel, Bioconductor’s unified interface for parallel processing in R (Morgan et al., n.d.). Additionally, MotifPeeker leverages ENCODE (Kheradpour and Kellis, 2014) and JASPAR (Rauluseviciute et al., 2024) APIs to automatically fetch alignment, peak and motif files if the user supplies the corresponding identifiers.

### Usage

Users can install MotifPeeker on Unix and Mac systems via BiocManager or GitHub remotes, provided the latest MEME Suite is installed. Alternatively, MotifPeeker is available through Docker and Singularity using containers hosted on the GitHub Container Registry, eliminating the need to manually install dependencies and the MEME Suite. The package can then be loaded up in R to run the main *MotifPeeker* function for the analysis. The function accepts multiple inputs, with the primary ones being peak_files, a vector specifying either paths to peak files for comparison or their ENCODE IDs for automatic download, and genome_build, which specifies the reference genome. Users can provide a character string such as “hg19”, “hg38”, “mm10”, “mm39”, or supply a *BSgenome* object from Bioconductor for any other genome build. Similar to *peak_files*, users can also pass vectors of *alignment_files* to compute additional general statistics like the fragment of reads in peaks (FRiP) score. To customise the labels and experiment classification of each input dataset, the *exp_labels* and *exp_type* parameters can be used. Motif files or JASPAR identifiers for motifs relevant to the target protein can be supplied via the motif_files parameter to enable reporting of summit-to-motif distances and known-motif enrichment analysis. The parameter accepts motifs in popular formats such as JASPAR, MEME, and TRANSFAC, or alternatively, users may pass a u*niversalmotif* object to utilise custom motif formats (Tremblay, 2024). Additional run customisation is also available in the form of other parameters, such as the number of motifs to find (*motif_discovery_count*) during the motif discovery step and the minimum number of consecutive nucleotide repeats required to filter out a discovered motif from the analysis (*filter_n*). A full list of function parameters, along with their usage guidance, is available here: https://neurogenomics.github.io/MotifPeeker/reference/MotifPeeker.html

Upon completion of the run, the HTML report is automatically launched in the user’s default web browser. If the user opts to save the run files (*save_runfiles=TRUE*), the intermediate output generated by the MEME Suite tools are also saved to the output folder. As another option, the HTML report includes embedded download buttons for accessing output files from each step, as well as files for segregated peaks. Computationally intensive steps, such as motif discovery, can be accelerated by specifying an appropriate BiocParallel backend using the *BPPARAM* parameter, ensuring that the chosen backend aligns with the available computational resources. Users can also use the exported core functions from MotifPeeker (such as *read_peak_file, read_motif_file, motif_enrichment, summit_to_motif, find_motifs*) into their custom analysis pipelines. Comprehensive documentation for all exported functions is available at https://neurogenomics.github.io/MotifPeeker/reference/index.html

### Output

The HTML report is organised into three, each including brief descriptions, interactive plots, and tables. (The first section focuses on general statistics computed for all alignment and peak files to provide context for comparisons. These include the fraction of reads in peaks (FRiP) score, which quantifies the proportion of mapped reads falling within peak regions, and peak width distributions, where the average and median peak widths are calculated. Additionally, summit to motif distances are assessed by plotting the distribution for each dataset and each supplied motif, together with the mean absolute distance which shows how far the detected signal lies from the motif occurrence. Users may also enable bootstrapping to estimate the distribution of these distances as the peaks are resampled. The summit refers to the nucleotide position with the highest fragment pileup, as determined by MACS. The second section covers known motif enrichment analysis, which computes and plots motif enrichment counts and percentages for segregated peaks in each comparison. The third section focuses on discovered motif enrichment analysis, where motif similarity scores are plotted using the Pearson Correlation Coefficient (PCC) across different segregated peak comparisons. Descriptions for each heatmap are provided in Table 1. For each discovered motif, the closest known motif in the JASPAR database is identified and presented in a table.

**Table 1.** Summary of motif similarity heatmaps for the discovered motif enrichment analysis section.

| Heatmap Label | Description |
| --- | --- |
| Common Motif Comparison | High correlation between motifs validates that shared peaks in reference and comparison datasets are enriched for similar motifs, suggesting consistent binding of shared regulatory factors across the experiments. |
| Unique Motif Comparison | This plot examines whether unique peaks from both experiments capture similar motifs. Highly correlated motifs indicates that comparisons miss out on capturing peaks with a likely legitimate motif. This may be caused by strict peak-calling thresholds, which potentially exclude true positive signals. |
| Cross Motif Comparisons A/B | These plots compare motifs from unique peaks in one experiment to motifs from common peaks in the other experiment (refer to axis labels). High motif correlation suggests that the dataset contributing unique peaks exhibits greater sensitivity, effectively capturing a larger proportion of motifs shared between the two experiments. |

We demonstrate our package using ChIP-Seq alignment files from ENCODE (Kheradpour and Kellis, 2014) with accession IDs ENCFF172XLC, ENCFF411OOS and ENCFF091ODJ, along with a TIP-Seq alignment file generated by (Bartlett et al., 2021). Peak files were subsequently generated using the MACS3 peak caller (Zhang et al., 2008). All experiments were conducted on HCT116 colorectal adenocarcinoma cells, targeting the CCCTC-binding factor (CTCF). For known motif enrichment analysis, motif files for CTCF and its structurally similar counterpart, CTCF-Like (CTCFL), were retrieved from the JASPAR database with matrix IDs MA1930.2 and MA1102.3, respectively. Several plots from the report are presented in Figure 2, and the full report can be viewed by selecting “CTCF ChIP-Seq – TIP-Seq” in the “Example Reports” vignette (https://neurogenomics.github.io/MotifPeeker/articles/examples.html).

**Figure 2.**
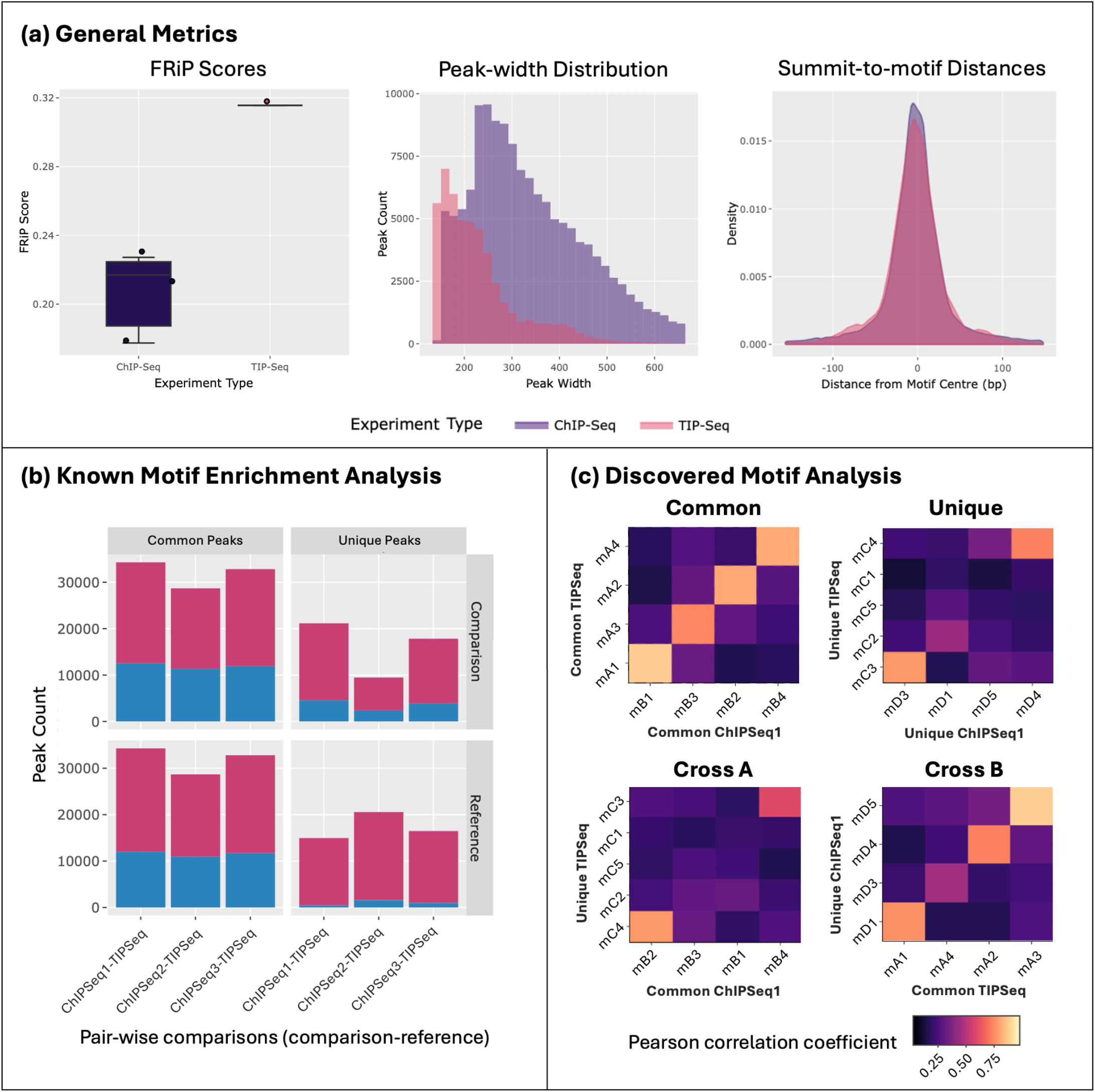
Comparative analysis of ChIP-Seq and TIP-Seq performance in profiling CTCF binding sites in HCT116 colorectal adenocarcinoma cells shows that while TIP-Seq produces narrower peaks with less background, it misses out on capturing genuine signals. (a) TIP-Seq exhibits lower noise (high FRiP score), improved precision (smaller peak widths) and comparable accuracy (similar summit-to-motif distances) when compared to ChIP-Seq. FRiP scores, peak-width distribution, and summit-to-motif distances for peaks grouped by the experimental method are plotted in order. (b) A higher proportion of peaks unique to ChIP-Seq contain the expected CTCF motif compared to the peaks unique to TIP-Seq (relative to ChIP-Seq). The bar graph shows the counts of enriched (blue) and non-enriched (red) peaks for the CTCF motif MA1930.2 when comparing the reference TIP-Seq dataset against three other comparison ChIP-Seq datasets. (c) While ChIP-Seq continues to capture motifs similar to those found in the common peak groups, TIP-Seq identifies additional unique motifs; however, these motifs appear irrelevant to the target CTCF motif, suggesting limited specificity in certain cases. The heatmaps show the Pearson correlation coefficients (PCC) between motifs discovered in the common and unique peak sets from the TIPSeq and ChIPSeq1 datasets, indicating the level of overlap and similarity in the top motifs found for each set. The “Common” heatmaps show correlations between motifs found in overlapping peak regions, while the “Unique” heatmaps compare motifs from peaks specific to each method. The high correlations observed in the “Cross B” heatmap, which compares motifs from unique ChIP-Seq peaks to common TIP-Seq peaks, suggest that TIP-Seq may underrepresent some legitimate regulatory motifs captured by the more sensitive ChIP-Seq approach. Conversely, the “Cross A” heatmap, comparing motifs from unique TIP-Seq peaks to common ChIP-Seq peaks, indicates that TIP-Seq can identify some distinct regulatory elements not detected by the ChIP-Seq method. Discovered motifs are labelled on the axes using identifiers with prefixes mA, mB, mC and mD, which stand for motifs discovered in common TIPSeq, common ChIPSeq1, unique TIPSeq and unique ChIPSeq1 peak sets, respectively.

## Conclusion

We developed MotifPeeker, an open-source R package that uses motif enrichment as a novel metric for comparing and benchmarking epigenomic datasets. Available through Bioconductor and GitHub, the package generates an HTML report with interactive plots and detailed descriptions in a single function call, making the analysis accessible even to users with limited computational experience. The results enable researchers to identify and validate epigenomic profiling techniques while also highlighting potential improvements in experimental protocols and data processing strategies. MotifPeeker is an actively maintained software tool that continues to see ongoing improvements, with plans to expand the supported peak file formats and genome builds.

## Acknowledgements

We sincerely thank Sarah Marzi and Alexi Nott for their valuable contributions to the discussions on early findings and for their insightful suggestions that helped shape new functionalities in MotifPeeker during its development. This paper was typeset with the bioRxiv word template by @Chrelli: www.github.com/chrelli/bioRxiv-word-template

